# Age-related differences in structural and functional prefrontal networks during a logical reasoning task

**DOI:** 10.1101/854224

**Authors:** Maryam Ziaei, Mohammad Reza Bonyadi, David C. Reutens

## Abstract

In logical reasoning, difficulties in inhibition of currently-held beliefs may lead to unwarranted conclusions, known as belief bias. Aging is associated with difficulties in inhibitory control, which may lead to deficits in inhibition of currently-held beliefs. No study to date, however, has investigated the underlying neural substrates of age-related differences in logical reasoning and the impact of belief load. The aim of the present study was to delineate age differences in brain activity during a syllogistic logical reasoning task while the believability load of logical inferences was manipulated. Twenty-nine, healthy, younger and thirty, healthy, older adults (males and females) completed a functional magnetic resonance imaging experiment in which they were asked to determine the logical validity of conclusions. Unlike younger adults, older adults engaged a large-scale network including anterior cingulate cortex (ACC) and inferior frontal gyrus (IFG) during conclusion stage. Our functional connectivity results suggest that while older adults engaged the ACC network to overcome their intuitive responses for believable inferences, the IFG network contributed to higher control over responses during both believable and unbelievable conditions. Our functional results were further supported by structure-function-behavior analyses indicating the importance of cingulum bundle and uncinate fasciculus integrity in rejection of believable statements. These novel findings lend evidence for age-related differences in belief bias, with potentially important implications for decision making where currently-held beliefs and given assumptions are in conflict.

## Introduction

Logical reasoning requires drawing a necessary conclusion from given assumptions. Beliefs and prior knowledge, however, may overshadow these assumptions and lead to unwarranted conclusions; a phenomenon known as belief bias (De Neys, 2012; Tsujii et al., 2010). A number of factors influence belief bias including time constraints (Evans & Curtis-Holmes, 2005; Trippas et al., 2013), emotional content of the reasoning statements (Goel & Vartanian, 2011), and higher cognitive ability (Sá et al., 1999; Trippas et al., 2015). Besides, the ability to inhibit current beliefs (De Neys, 2012; Houde & Borst, 2015; Houde et al., 2000) is critical for adaptive behaviors (Houde et al., 2000) and arriving at a logical conclusion (Houde & Borst, 2015). Considering the critical role of logical reasoning in personal, complex political, and societal decisions, a reduced ability in inhibition of beliefs may indicate underlying neural changes, which could have consequences for daily decision-making.

Inhibitory control – a foundation of many cognitive functions such as attention and memory (Paxton et al., 2008) – shows age-related changes; improves from childhood to early adulthood but declines in later life. Age-related decline in inhibitory control may lead to difficulties in the inhibition of currently-held beliefs and, consequently, result in reduced logically correct decisions. Further, evidence suggests that normal aging is associated with a reduction of gray matter volume in frontal regions (Mann et al., 2011) and reduction of structural integrity of white matter tracts between frontal and other parts of the brain (Hasan et al., 2009; Treit et al., 2014; Wolf et al., 2014). Thus, understanding age-related differences in structural and functional networks involved in inhibitory control provides further insights to reasoning and decision-making in late adulthood.

To our knowledge, only two studies have investigated age-related differences in logical reasoning so far. De Neys and Van Gelder (2009) examined the relationship between deductive reasoning performance and cognitive control ability behaviorally and found that reasoning performance decreased among older adults when beliefs and logic were in conflict. No difference in performance was found when logic and beliefs agreed; belief inhibition was not required. In another study, Tsujii and colleagues (2010) examined the role of the inferior frontal gyrus (IFG) – a critical region in reasoning and inhibitory control – in the inhibition of current beliefs during a logical reasoning task using near-infrared spectroscopy. Unlike younger adults, older adults recruited the bilateral inferior frontal gyrus – when belief and logic were in conflict – suggesting a possible mechanism to compensate for age-related decline in inhibitory control. Although Tsujii et al’s (2010) finding was critical in highlighting the role of IFG in logical reasoning in aging, no study to date has investigated the underlying functional networks involved in logical reasoning and the impact of belief load in late adulthood.

In this study, we aimed to identify age-related differences in both structural and functional substrates of logical reasoning when the belief load of inferences and currently-held beliefs are either in conflict or agreement. Younger and older adults completed a syllogistic reasoning task in which they identified if a conclusion logically followed given assumptions. The believability content of the inferences was either congruent (believable; all parrots are birds) or incongruent (unbelievable; all lizards are mammals) with currently-held beliefs. In this study, in order to measure cognitive control, we define *performance* as accepting unbelievable or rejecting believable statements. Given that more cognitive control is required to suppress currently-held beliefs, we anticipated that cognitive control areas, such as anterior cingulate cortex (ACC) and inferior frontal gyrus, would contribute to higher performance (correctly rejecting believable and correctly accepting unbelievable statements). Converging evidence highlights the importance of IFG in inhibitory control (Brass et al., 2005) and the resolution of belief-logic conflict in logical reasoning (Prado et al., 2011; Reverberi et al., 2012; Reverberi et al., 2010; Stollstorff et al., 2012), supporting the choice of IFG as a seed for functional connectivity analyses. Furthermore, mounting evidence supports the critical role of ACC in self-regulation functions such as monitoring of conflict (Botvinick et al., 2001; Botvinick et al., 2004), regulating emotions (Buhle et al., 2014; Bush et al., 2000; Ziaei et al., 2017), conflict monitoring (Vassena et al., 2017), and responding to errors (Holroyd & Coles, 2002), all of which are essential for inhibiting currently-held beliefs during logical reasoning. The role of ACC was also highlighted in deductive reasoning in the absence of any visual input (Knauff et al., 2002).

Using functional connectivity analysis, we investigated functional networks of the ACC and IFG during a logical reasoning task. Existing theories argue that unbelievable inferences encourage a more thorough reasoning because of the conflict between belief load and logic. According to the mental model theory, individuals construct mental models from syllogisms and if the syllogisms’ belief load is in conflict with the mental model, it initiates a search for an alternative model of the premises (Johnson-Laird, 2001; Johnson-Laird, 2010; Johnson-Laird et al., 2015). Therefore, *accepting* unbelievable statements and *rejecting* believable ones prompts conflict and thus demands higher cognitive control. Given the well-documented inhibitory control deficits in late adulthood (Gazzaley & D’Esposito, 2007; Hasher & Zacks, 1988), we hypothesized that older adults would show difficulties to accept an unbelievable statement or reject a believable one correctly, both of which require higher cognitive control ability.

Given the existing evidence on the relationship between the lower white matter integrity and age-related cognitive decline (Madden et al., 2009), we examined whether the structural strength of pathways connecting frontal and other brain areas contribute to correct logical reasoning and if this relationship would differ between younger and older adults. We selected the cingulum bundle and uncinate fasciculus tracts that are mainly involved in inhibitory control (Li et al., 2018), executive functioning (Grieve et al., 2007), and belief updating (Moutsiana et al., 2015). In line with our functional connectivity predictions, we anticipated rejecting believable and accepting unbelievable statements correctly would require higher cognitive control, possibly relying on the white matter integrity. Furthermore, it is expected that the integrity of white matter tracts underlies the relationship between age and logical reasoning performance. That is, older adults with greater white matter integrity would be more capable in correctly rejecting believable statements and correctly accepting unbelievable ones.

## Material and Methods

### Participants

Thirty-two, healthy, older and thirty-one, healthy, younger adults participated in this study. Two older and two younger adults were excluded from the analysis due to brain signal loss and extensive movement, leaving 30 older adults (aged 61-78 years; *M* = 70.34, *SD* = 4.27; 15 females) and 29 younger adults (aged 18-26 years; *M* = 21.13, *SD* = 2.72; 15 females). All younger adults were undergraduate students at the University of Queensland who were reimbursed either with course credits or $15 AUD per hour. Older adults were volunteers from the community recruited through advertising in public notice boards of different clubs, libraries, churches, and the University of Queensland’s Aging Mind Initiative. Older adults were reimbursed $20 AUD per hour. Before taking part in this study, all participants were screened for MRI compatibility as well as claustrophobia, mood disorders such as depression and anxiety, neurological and psychiatric medication including cardiovascular and epilepsy. All participants were right-handed, English speakers who had normal or corrected-to-normal vision using MRI compatible glasses with no history of psychiatric illnesses, head or heart surgery, and neurological impairment. Both age groups were similar in education years and gender. Older adults underwent screening to rule out cognitive decline on a widely used dementia screen, the Mini Mental State Examination (Folstein et al., 1975). All older adults scored above the recommended cut-off of 27 (*M* = 29.34, *SD* = 0.82). Participants took part in two separate sessions of testing, the first involving fMRI scanning and the second involved behavioral/neuropsychological assessment. Informed consent was obtained from all individuals included in the study and they were debriefed upon the completion of second session. All procedures performed in studies involving human participants were in accordance with the ethical standards of the institutional and/or national research committee and with the 1964 Helsinki declaration and its later amendments or comparable ethical standards. The study was approved by the independent Bellberry Human Research Ethics Committee.

### Task materials

A logical *statement* in this study follows a generic form of <*quantifier, subject, copula, predicate>*, e.g., All parrots are birds, where “All” is a quantifier, “parrots” is a subject, “are” is a copula, and “birds” is a predicate. There are three choices for the quantifier that are “All”, “No”, and “Some”, and two choices for the copula that are “are” and “are not”. Four propositions are generated by combining a quantifier and a copula: Proposition type *A* that is a combination of “All” and “are”; Proposition type *E* that is a combination of “No” and “are”; Proposition type *I* that is a combination of “Some” and “are”; Proposition type *O* that is a combination of “Some” and “are not”. Other combinations (e.g., “All” and “are not”) were not used in this study.

A logical argument in this study included three statements, two premises and one conclusion, in the form of a standard syllogism (Table 1). In a premise, the subject and the predicate are arbitrary sets (e.g., dogs, mammals, furniture). Exactly one set is *shared* between two premises that may appear in either the subject or the predicate in either of the premises (*Set*_2_ in Table 1). Hence, an argument with two premises involves exactly three sets (*Set*_1_, *Set*_2_, and *Set*_3_), two uniquely used in each premise and one used in both. The conclusion of a syllogism is a statement about the sets that appear uniquely in premises (*Set*_1_ and *Set*_3_). A conclusion “follows” from the premises if the premises provide conclusive evidence to support it. Otherwise, the conclusion “does not follow” from the premises. This includes the conclusion that is wrong, given the premises, or is not completely supported by the premises.

**Table 1.** Form of syllogisms, quantifiers, subjects, copula, and predicate used in the experimental design.

|  | Quantifier | Subject | Copula | Predicate |
| --- | --- | --- | --- | --- |
| Premise 1 | All | Set<sub>1</sub> | is | Set<sub>2</sub> |
| Premise 2 | No | Set<sub>2</sub> | is | Set<sub>3</sub> |
| Conclusion | Some | Set<sub>3</sub> | is not | Set<sub>1</sub> |

Conclusions and premises can be believable or unbelievable statements. For example, “all birds are mammals” is an unbelievable statement while “all parrots are birds” is a believable statement. In addition, premises may provide a neutral context, for example, “all parrots are nickhomes”, where “nickhomes” is a meaningless pseudo-word. This type of statement was only used in the premises in such a way that the pseudo-word was always the shared set between the premises. Hence, the premises were either believable, unbelievable, or neutral while the conclusion was either believable or unbelievable. The phrases “believable”, “neutral”, and “unbelievable” mean that the first premise was always a believable statement while the second was either believable, neutral, or unbelievable, respectively. The following are two examples of syllogisms with a believable premise/unbelievable conclusion and an unbelievable premise/believable conclusion, respectively:

All sparrows are birds; all birds are animals; therefore, no animals are sparrows: believable premise/unbelievable conclusion

All oranges are citrus; no citrus are fruits; therefore, some fruits are not oranges: unbelievable premise/believable conclusion

We used propositions *A* (All, are) and/or *E* (No, are) in both of premises (counter balanced across all runs). We avoided the use of proposition *E* in both premises of the same argument as it simplifies the reasoning task, making the premises AA, AE, and EA. For the conclusions, however, we used the propositions required to ensure that the conclusions that followed the premises and those that did not follow the premises were balanced.

We also balanced the way the sets in the subject and predicate of the premises were ordered. In this experiment, we used two types of ordering: i) the shared set belonged to the subject of premise 1 and the subject of premise 2, and ii) the shared set belonged to the predicate of premise 1 and the subject of premise 2. The conclusion takes the unique sets from each premise, combines them through a proposition, and the logical reasoning task is to decide if the generated statement follows from the premises or not. Whether the subject of the conclusion was drawn from the first or second premise was also counterbalanced in each condition.

An algorithm was developed that creates all the syllogisms based on criteria specified in Table 1. A total of 96 syllogisms was generated, within six conditions: 1. believable premise/believable conclusion, 2. believable premise/unbelievable conclusion, 3. unbelievable premise/believable conclusion, 4. unbelievable premise/unbelievable conclusion, 5. neutral premise/believable conclusion, 6. neutral premise/unbelievable conclusion (see Supplementary Material for access to all of the syllogisms used in the study).

### Experimental design

During the logical reasoning task, participants ascertained if the conclusion statement logically followed from the premises. They responded with two keys on an MRI-compatible response box. The first premise was presented for 2 seconds followed by a second premise for 4 seconds. After the second premise, the conclusion statement was presented for 12 seconds (Figure 1). To minimize the working memory load, all statements remained on the screen until the end of the presentation of the conclusion. A fixation cross was presented after the conclusion, which was randomly jittered using four-time intervals across all runs: 0.5 seconds (24 trials), 1 second (24 trials), 1.5 seconds (24 trials), and 2 seconds (24 trials). The task consisted of 6 runs, each run lasting for 5.16 minutes. Overall, 96 syllogisms were presented across all runs with 16 syllogisms per each condition.

**Figure 1.**
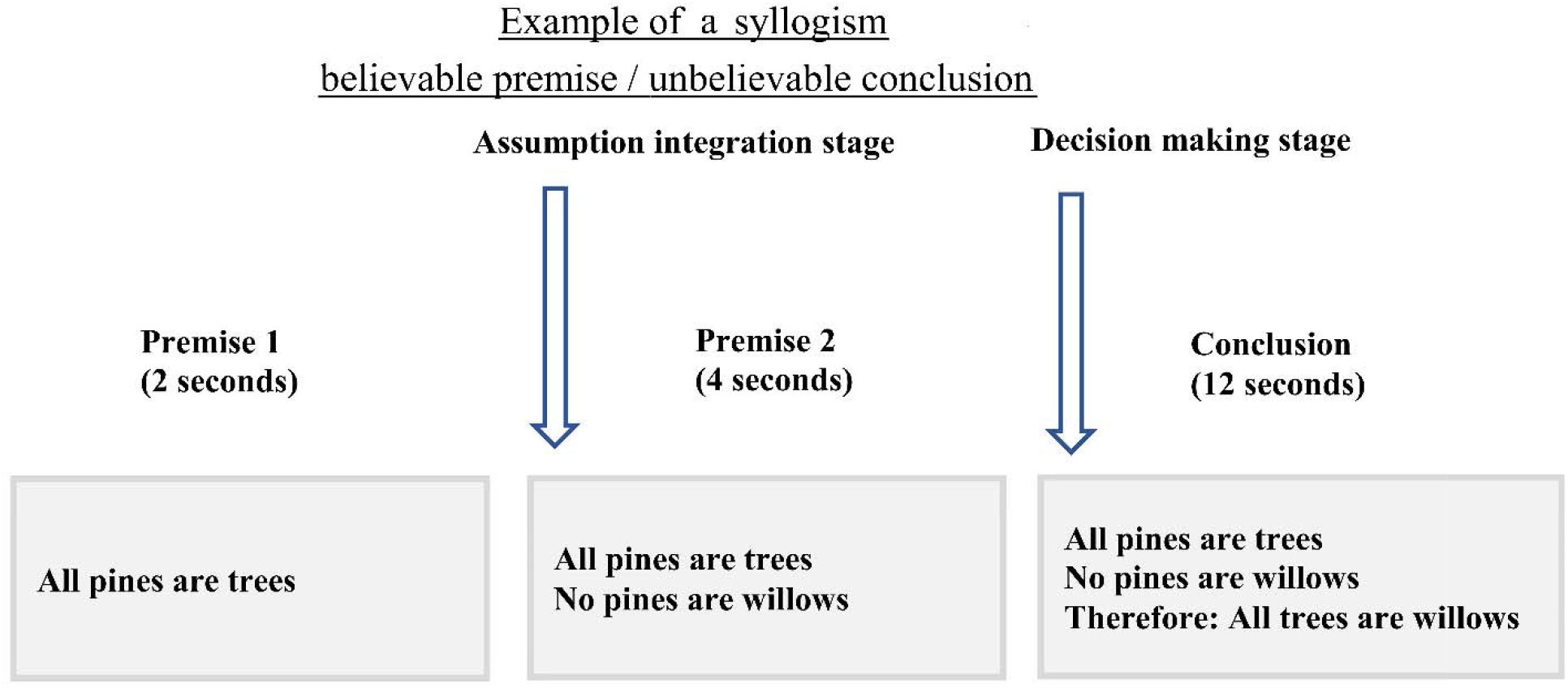
The timing and example of the experimental design. In this experiment, participants were presented with the first premise for two seconds followed by the second premise that was shown for 4 seconds. During the conclusion stage, 12 seconds, participants were asked to choose if the conclusion followed the two premises or not, responding with an MRI compatible response box. All statements remained on the screen to lower working memory load required to perform the task.

### Procedure

The experiment consisted of two sessions: scanning session and behavioral/neuropsychological assessment session. Each participant’s scanning session lasted 45 minutes and included two components: structural MRI and functional MRI with the logical reasoning task. Prior to the scan, participants were verbally instructed about the task and a practice run was continued until they were familiar with the timing and instruction of the task. Three runs of the task were presented before and three after the structural scans. Following completion of the scanning session, participants were asked to complete number of background measures.

### Background measures

During the behavioral/neuropsychological assessment session, all participants completed a number of tasks to assess executive functioning measured by Stroop task (Jensen & Rohwer, 1966) and Trail Making Test (Reitan & Wolfson, 1986); emotional well-being measured by Depression, Anxiety, Stress Scale (DASS-21, Lovibond & Lovibond, 1995); and intelligence measured by the National Adult Reading Test (Nelson, 1982). Descriptive and inferential statistics of background measures are reported in Table 2.

**Table 2.**
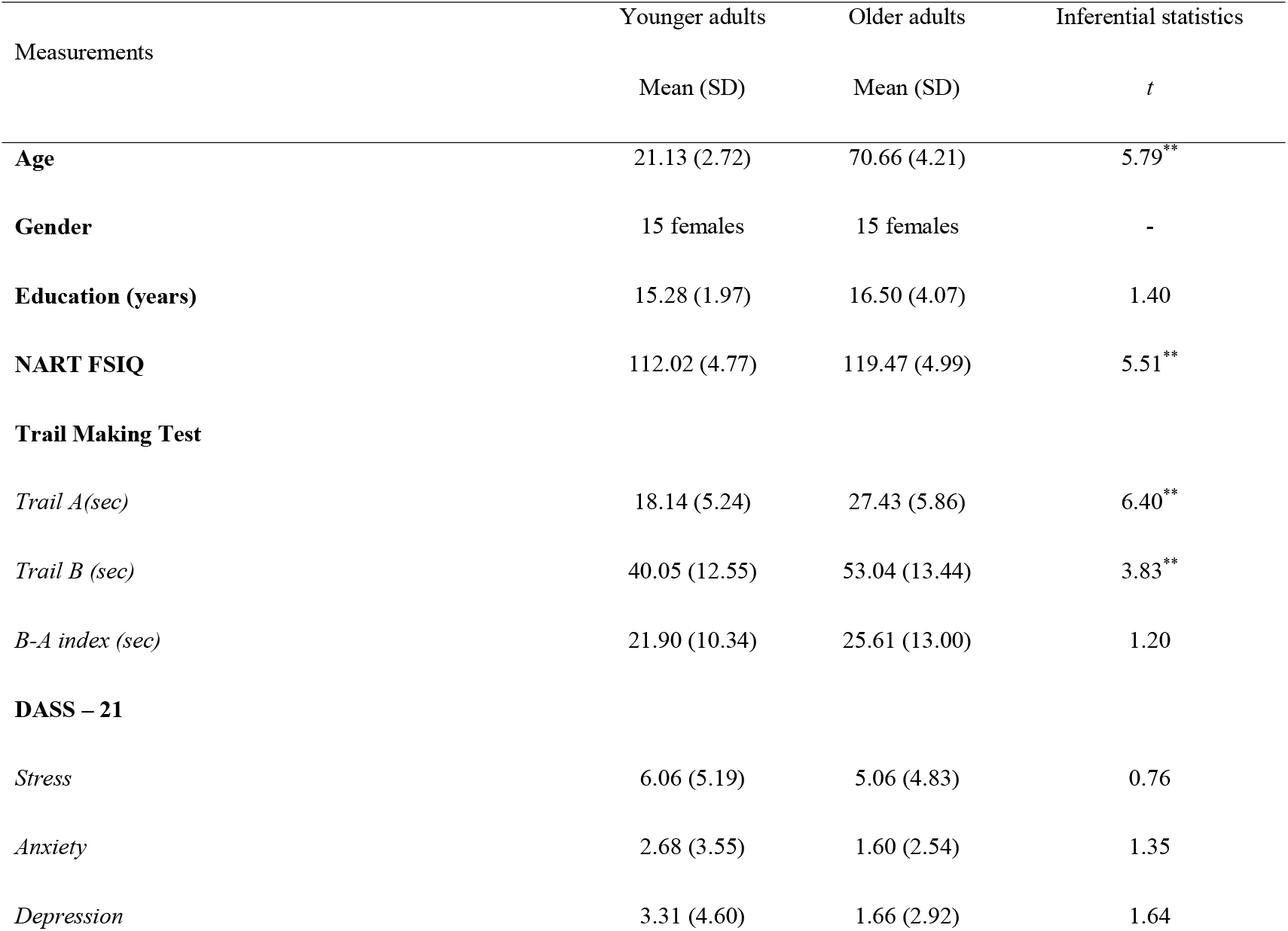
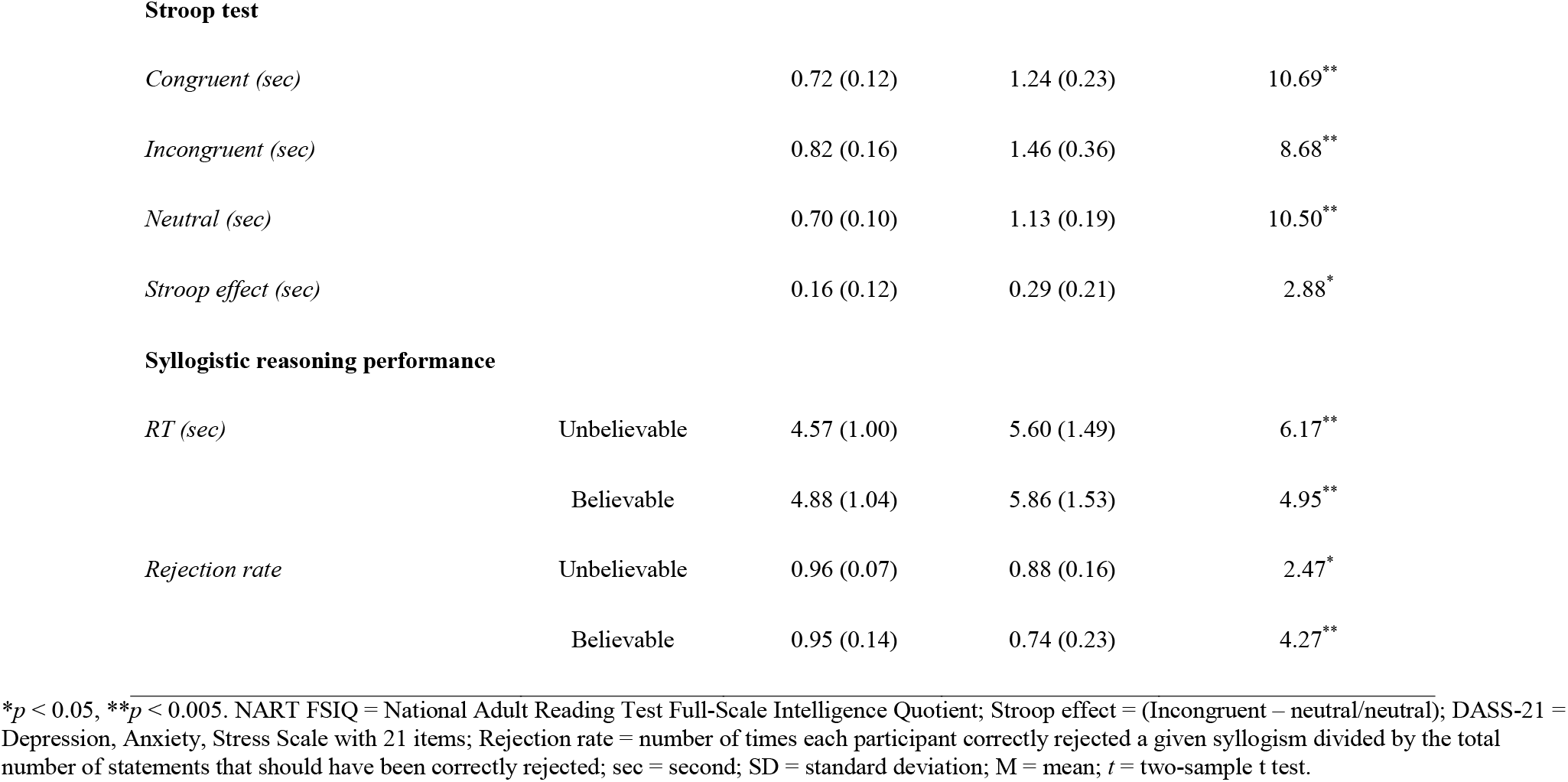
Descriptive and inferential statistics of performance on background measures and logical reasoning task.

|  | Younger adults | Older adults | Inferential statistics |
| --- | --- | --- | --- |
| Measurements | Mean (SD) | Mean (SD) | *t* |
| **Age** | 21.13 (2.72) | 70.66 (4.21) | 5.79** |
| **Gender** | 15 females | 15 females | - |
| **Education (years)** | 15.28 (1.97) | 16.50 (4.07) | 1.40 |
| **NART FSIQ** | 112.02 (4.77) | 119.47 (4.99) | 5.51** |
| **Trail Making Test** |  |  |  |
| *Trail A(sec)* | 18.14 (5.24) | 27.43 (5.86) | 6.40** |
| *Trail B (sec)* | 40.05 (12.55) | 53.04 (13.44) | 3.83** |
| *B-A index (sec)* | 21.90 (10.34) | 25.61 (13.00) | 1.20 |
| **DASS – 21** |  |  |  |
| *Stress* | 6.06 (5.19) | 5.06 (4.83) | 0.76 |
| *Anxiety* | 2.68 (3.55) | 1.60 (2.54) | 1.35 |
| *Depression* | 3.31 (4.60) | 1.66 (2.92) | 1.64 |

Stroop test
|  |  |  |  |  |
| --- | --- | --- | --- | --- |
| *Congruent (sec)* |  | 0.72 (0.12) | 1.24 (0.23) | 10.69** |
| *Incongruent (sec)* |  | 0.82 (0.16) | 1.46 (0.36) | 8.68** |
| *Neutral (sec)* |  | 0.70 (0.10) | 1.13 (0.19) | 10.50** |
| *Stroop effect (sec)* |  | 0.16 (0.12) | 0.29 (0.21) | 2.88* |

Syllogistic reasoning performance
|  |  |  |  |  |
| --- | --- | --- | --- | --- |
| *RT (sec)* | Unbelievable | 4.57 (1.00) | 5.60 (1.49) | 6.17** |
|  | Believable | 4.88 (1.04) | 5.86 (1.53) | 4.95** |
| *Rejection rate* | Unbelievable | 0.96 (0.07) | 0.88 (0.16) | 2.47* |
|  | Believable | 0.95 (0.14) | 0.74 (0.23) | 4.27** |

\* $p < 0.05$ , \*\* $p < 0.005$ . NART FSIQ = National Adult Reading Test Full-Scale Intelligence Quotient; Stroop effect = (Incongruent – neutral/neutral); DASS-21 = Depression, Anxiety, Stress Scale with 21 items; Rejection rate = number of times each participant correctly rejected a given syllogism divided by the total number of statements that should have been correctly rejected; sec = second; SD = standard deviation; M = mean; $t$ = two-sample $t$ test.

### fMRI image acquisition

Functional images were acquired at the Centre for Advanced Imaging using a 3T Siemens scanner with a 32-channel head coil. The functional images were obtained using a whole-head T2*-weighted multiband sequence (473 interleaved slices, repetition time (TR) = 655ms, echo time (TE) = 30ms, voxel size = 2.5mm^3^, field of view (FOV) = 190mm, flip angle = 60°, multi-band acceleration factor = 4). High-resolution T1-weighted images were acquired with an MP2RAGE sequence (176 slices with 1mm thickness, TR = 4000ms, TE = 2.89ms, TI = 700ms, voxel size = 1mm^3^, FOV = 256mm). To minimize the noise and head movement, participants were provided with cushions and earplugs around their head inside the head coil. Participants observed the task on a computer screen through a mirror mounted on top of the head coil.

#### Preprocessing

For functional analysis, T2*-weighted images were preprocessed with Statistical Parametric Mapping Software (SPM12; http://www.fil.ion.ucl.ac.uk/spm) implemented in MATLAB 2015b (Mathworks Inc., MA). Following realignment to a mean image for head-motion correction, images were segmented into gray and white matter. Then, images were spatially normalized into a standard stereotaxic space with a voxel size of 2 mm^3^, using the Montreal Neurological Institute (MNI) template, and then spatially smoothed with a 6 mm^3^ Gaussian Kernel. None of the participants included in the analysis had head movement exceeding 1mm.

#### fMRI analyses

The imaging data were analyzed using a model-free, multivariate analytical technique, Partial Least Squares (PLS; McIntosh et al., 1996; McIntosh et al., 2004). For a detailed tutorial and review of PLS, see Krishnan et al. (2011), as implemented in PLS software running on MATLAB 2012b (The MathWorks Inc., MA). In brief, PLS analysis uses singular value decomposition (SVD) of a single matrix that contains all data from all participants to find a set of orthogonal latent variables, which represent linear combinations of the original variables (latent variables, LV). Usually, the first latent variable accounts for the largest covariance of the data, with a progressively smaller amount of covariance for subsequent latent variables. Each latent variable delineates cohesive patterns of brain activity related to experimental conditions. Therefore, each latent variable consists of a singular image of voxel saliences (*i.e*., a spatiotemporal pattern of brain activity), a singular profile of task saliences (*i.e*., a set of weights that indicate how brain activity in the singular image is related to the experimental conditions, functional seeds, or behavioral/anatomical covariates), and a singular value (*i.e*., the amount of covariance accounted for by the latent variable). Rather than making assumptions about conditions or imposing contrasts for each pattern, PLS decomposes all images into a set of patterns that capture the greatest amount of covariance in the data. Therefore, PLS enables differentiation of the contribution of brain regions associated with task demands, functional seed activity, and behavioral or anatomical covariates. The reliability of the salience for each brain voxel was determined using 100 bootstrap resampling steps (Efron & Tibshirani, 1985). Peak voxels with a bootstrap ratio (*i.e*., salience/standard error) > 3.0 were considered to be reliable, as this approximates *p* < 0.01 (Sampson et al., 1989). Because PLS is a multivariate approach and considers all voxels simultaneously, it does not require correction for multiple comparisons (McIntosh et al., 2004), which remains a fraught issue for the GLM framework (Eklund et al., 2016).

PLS offers multiple advantages compared to other traditional methods such as general linear model (GLM) and thus, has been used as a primary method of analyzing fMRI data in this experiment. PLS is constrained to examine the covariance between brain activity and task design rather than explicitly modeling the hemodynamic response function (HRF). In this regard, this method enables examining robust patterns of activity only associated with the experimental conditions. Along the same lines, PLS is a multivariate approach, capable of analyzing multiple conditions simultaneously to model covariance of response across conditions. Due to this, PLS does not require correction for multiple comparisons, which is a fraught issue for the GLM framework. Regarding the power, PLS is more sensitive than the GLM, particularly when the dependent variables are correlated. Thus, by using PLS, limitations associated with age-related differences in BOLD and Hemodynamic Response Function modelling would not impact the results (Fynes-Clinton et al., 2019; Ziaei et al., 2016).

### DTI data acquisition

High-angular resolution Diffusion Weighted Images (DWI) were acquired using High Angular Resolution Diffusion Imaging (HARDI) sequence; 60 slices, TR = 8600 ms, TE = 109 ms, Diffusion direction = 64, b value = 3000 s/mm2, size = 2.3mm^3^) and processed using FSL software (Fischl, 2012).

#### DTI data processing

The DWI images were first corrected for the eddy current distortion and head motion using the FMRIB’s Diffusion Toolbox (FDT) (Andersson & Sotiropoulos, 2016). Non-brain tissues were removed from the corrected images using the Brain Extraction Tool (Smith, 2002). We then used the FDT to locally fit the diffusion tensor model at each voxel and prepare the map of fractional anisotropy (FA) values throughout the brain of each subject. Fractional anisotropy is a general marker of white matter integrity that reflects the coherence within a voxel and fiber density (Alexander et al., 2007; Beaulieu, 2002). In the absence of other diffusivity measures, FA is accepted as a nonspecific marker of microstructural change (Alexander et al., 2007). The FA map of each subject was then realigned to match the standard brain template, MNI 152 T1 1mm space, using the FMRIB’s Linear Image Registration Tool (FLIRT) (Greve & Fischl, 2009; Jenkinson et al., 2002; Jenkinson & Smith, 2001), with 12 degrees of freedom and trilinear interpolation. This provides an affine transformation that ensures the transformed FA maps are in the same 3D coordinate system as of one another. We then generated the desired white matter mask for the tract of interest using the ICBM-DTI-81 white-matter atlas (Hua et al., 2008; Oishi et al., 2010; Wakana et al., 2007); We generated the masks for inferior Longitudinal Fasciculus, Cingulum, and Uncinate Fasciculus tracts that have been shown to play a role in executive functioning, inhibitory control, and are particularly impacted in aging (Hasan et al., 2009; Li et al., 2018). The average of FA values in the tract of interest for each subject were then calculated and used in structure-function and structure-behavior analyses.

### Whole-brain task-related analysis

We examined how belief load of the conclusions influenced the logical reasoning in the whole-brain analysis (task PLS). This analysis focused on the time during the task that participants had to decide about the logical validity of conclusions, hence, referred to as decision-making stage from now on. Onsets from the beginning of the *conclusions* for all six experimental conditions were included in this analysis aiming to identify the brain activation related to the conclusions’ belief load. However, to ensure that brain activities reported are related to the reasoning and are not confounded with other processes that are involved during reasoning (visual and language processes, for instance), brain activities from the 3^rd^ to 9^th^ TRs (correspond to 1.96 - 5.89 seconds after presentation of the conclusion statement) are reported here. This time interval in the conclusion contains time bins with maximum response probability (see Supplementary Results for distribution of responses).

### Brain-behavior connectivity analyses

The aim of brain-behavior connectivity analysis was to examine whether the functional connectivity of the frontal regions differs between younger and older adults during the correct rejection of believable and correct acceptance of unbelievable inferences. Furthermore, we investigated whether there is any association between the task-related brain functional networks of frontal regions with behavioral performance. Given that the whole-brain activation pattern during conclusion was stronger for older than younger adults, to reduce the biases toward one age group, we have chosen two seed regions from the meta analyses reported in Neurosynth (Yarkoni et al., 2011) and therefore, included both age groups in the brain-behavior connectivity analyses. For functional connectivity, inferior frontal gyrus (IFG: Right: [48 22 8]; Left: [-48 22 16]) and anterior cingulate cortex (ACC: [0 32 22]) were chosen. The selection of these two seeds was primarily based on previous studies highlighting the importance of these regions in deductive reasoning (Prado et al., 2011; Reverberi et al., 2012; Reverberi et al., 2010; Stollstorff et al., 2012) as well as other underlying cognitive processes that are crucial for inhibiting beliefs such as monitoring conflicts and regulating emotions.

We conducted two separate analyses to identify correlation between activity of the anterior cingulate cortex and IFG with activity in the rest of the brain and behavior in both age groups. Our behavioral measure was rejection rate averaged across all trials in each experimental condition. In this study, we use the onsets of conclusion statements for the decision-making stage using an event-related design analysis.

The procedure of brain-behavior functional connectivity analyses involved extracting blood-oxygen-level dependent (BOLD) values form each of the selected seeds, ACC and IFG, from the onset of conclusion stage. The activity for each seed as well as the mean of rejection rate for each condition were then correlated with the activity in the rest of the brain within each experimental condition, across both age groups. This forms a covariance matrix which then is decomposed with singular value decomposition (SVD), resulting in a set of independent, orthogonal latent variables (LV). For each LV, “brain scores” indicate the degree to which each participant shows brain activity pattern across experimental conditions. Brain scores, which are calculated for each participant, were then correlated with the ACC and IFG BOLD values to examine the relationship between whole-brain brain activity pattern and activity in these two seeds. The results are displayed with the Pearson product-moment correlation coefficient between brain scores from whole-brain and other variables (rejection rate, and ACC/IFG percent signal change) for each experimental condition for each age group. We have also conducted additional analyses using endorsement rate as a behavioral measure that are reported in the Supplementary Results.

### Structure-function-behavior analyses

To investigate the role of structural integrity underlying logical reasoning, we chose three tracts: inferior Longitudinal Fasciculus, Cingulum, and Uncinate Fasciculus. These tracts include several critical frontal areas such as the anterior cingulate and orbitofrontal cortex, regions that were consistently found in our whole-brain task-related functional findings during the conclusion stage (Table 3) and temporal areas. For instance, the altered integrity of the Uncinate Fasciculus tract due to age (Meltzer-Baddeley et al. 2011) may reduce control on regions associated with processing semantic knowledge. We also need to note that inferior Longitudinal Fasciculus tract was chosen as a control to demonstrate the specificity of the effect to the regions such as cingulate and temporal lobe rather than general white matter integrity. To assess whether structural integrity of selected white matter tracts was related to the brain activation during the task and behavioral performance, we performed PLS analyses where the structural integrity of the cingulum bundle, uncinated fasciculus, and inferior longitudinal fasciculus were correlated with the task activation for both age groups as well as behavioral performance, rejection rate. The structural integrity (FA values) in each tract, both left and right sides, were included and correlated with the task-related functional activation during. For these analyses, brain scores, which are calculated for each participant, were correlated with the FA values of these tracts to examine the relationship between whole-brain activity pattern, structural integrity of white matter tracts, and behavioral performance. All experimental conditions, except neutral ones, were included in the analyses.

**Table 3.** Whole-brain task-related results during the decision-making stage.

| Regions | Hem | MNI coordinates | BSR |
| --- | --- | --- | --- |
|  |  | XYZ |  |
| Anterior cingulate cortex | R | 2 54 10 | 7.9374 |
|  | R | 0 46 -4 | 8.3378 |
| Inferior frontal gyrus (p. Triangularis | L | -52 32 10 | 6.3213 |
| Middle frontal gyrus | L | -26 32 46 | 7.208 |
| Superior temporal gyrus | L | -58 -12 -4 | 7.3709 |
|  | R | 64 -18 -6 | 7.0866 |
| Temporal pole | L | -40 18 -18 | 4.4267 |
| Precentral gyrus | R | 38 -20 52 | 8.443 |
| Postcentral gyrus | L | -58 -6 28 | 5.0609 |
| Superior parietal lobe | L | -6 -36 48 | 8.6503 |
| Lingual gyrus | L | -14 -74 -8 | 5.5967 |
| Fusiform gyrus | R | 26 -30 -18 | 5.7254 |
| Superior occipital gyrus | L | -20 -86 36 | 6.0812 |
|  | R | 16 -86 34 | 4.8526 |
| Middle occipital gyrus | L | -50 -74 6 | 6.0241 |
| Inferior occipital gyrus | R | 42 -78 -10 | 5.2606 |
| Occipital lobe | R | 42 -70 6 | 7.4544 |
|  | L | -30 -84 2 | 5.7525 |
| Hippocampus | R | 22 -14 -14 | 5.124 |

Hem = hemisphere; BSR = bootstrap ratio; MNI = Montreal Neurological Institute. Onsets of syllogisms from decision making, conclusion stage, were included in the analysis. These regions were more engaged by older adults for all experimental conditions relative to younger counterparts.

### Structure-behavior analyses

We performed a nonparametric bootstrapping mediation analyses to test the mediation model of FA values in white matter tracts as mediators of the relationship between age (independent variable) and rejection rate (dependent variable). Mediation is significant if the 95% confidence intervals for the indirect effect do not include zero (Preacher & Hayes, 2004; Preacher et al., 2007). Results were calculated based on the 10,000 bootstrapped samples for both left and right sides of white matter tracts, independently.

### Statistical analyses for behavioral data

It must be noted that a high performance in this task is defined by a high ratio of correctly rejecting believable or correctly accepting unbelievable statements. Given that rejecting a believable or accepting an unbelievable statement requires higher control over currently-hold beliefs, these two measures are reflecting higher cognitive control abilities. In other words, participants who can inhibit their currently-held beliefs when necessary, are able to reject a believable or accept an unbelievable statement, correctly. The rejection rate was the number of times each participant correctly rejected a given syllogism divided by the total number of statements that should have been correctly rejected. Endorsement rate was the number of times each participant accepted a given syllogism divided by the total number of statements that should have been correctly accepted. Repeated measures ANOVA were performed on dependent variables such as rejection rate and endorsement rate for syllogisms with believable and unbelievable loads from conclusion stage.

## Results

### Behavioral results

#### Rejection rate

A 2 (conclusion belief load: believable or unbelievable) by 2 (age group: younger and older adults) repeated measure ANOVA on correct rejection rate revealed a significant main effect of conclusion belief load (*F*(1,57) = 10.35, *p* = 0.002, 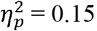), participants had higher rejection rate for unbelievable (*M* = 0.92, *SD* = 0.13) conclusions relative to believable (*M* = 0.84, *SD* = 0.22) conclusions (*t*(58) = 3.07, *p* = 0.003, *d* = 0.80). A main effect of age group reached a significant level (*F*(1,57) = 16.91, *p* = 0.000, 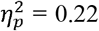), suggesting that older adults had lower performance relative to younger counterparts. An interaction between age group and conclusion conditions was also significant (*F*(1,57) = 8.32, *p* = 0.006, 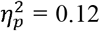), suggesting that the difference between two groups was much larger for the believable conclusion (*t*(57) = 2.48, *p* = 0.01, *d* = 0.65) than unbelievable ones (*t*(57) = 4.27, *p* = 0.00, *d* = 1.13).

#### Endorsement rate

A 2 (conclusion belief load: believable or unbelievable) by 2 (age group: younger and older adults) repeated measure ANOVA on endorsement rate revealed no significant results: main effect of conclusion belief load (*F*(1,57) = 0.24, *p* = 0.62, 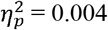), interaction between conclusion belief load and age group (*F*(1,57) = 3.33, *p* = 0.73, 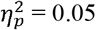), and main effect of age group (*F*(1,57) = 0.28, *p* = 0.59, 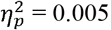).

### Imaging results

#### Whole-brain results

Our analyses during decision-making stage showed one significant latent variable which accounted for 49% of the variance of the data (*p* < 0.001). This latent variable consisted of a large-scale brain network including right ACC, bilateral IFG, bilateral superior temporal gyrus, left temporal pole, left superior parietal lobe, and right hippocampus (Table 3) that were more activated for all conditions among older adults relative to younger adults. Figure 2 shows brain activities based on different TRs from 3^rd^ to 9^th^ TRs.

**Figure 2.**
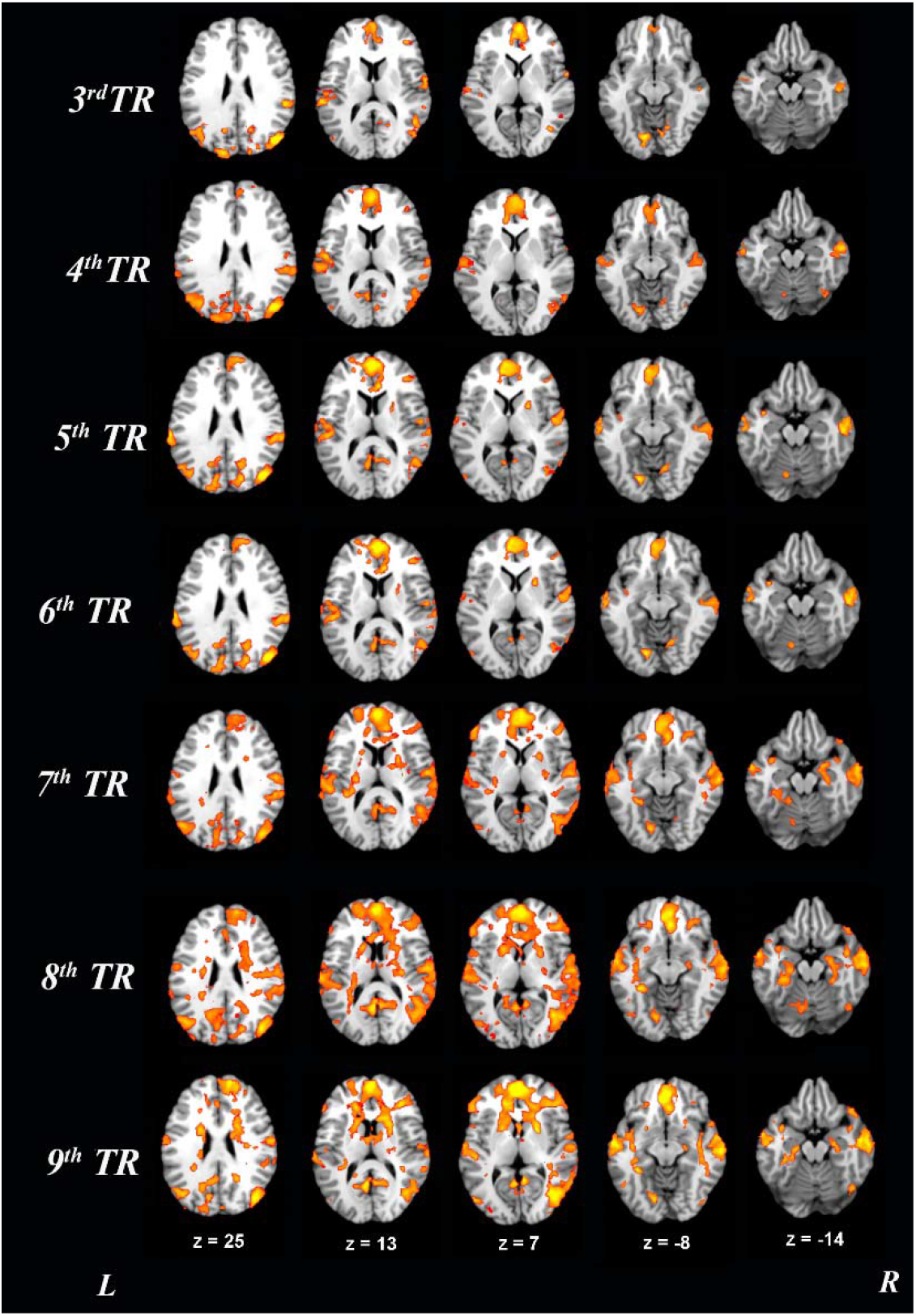
Whole-brain task-related results during the decision-making stage. Onsets of the conclusion stage for all six experimental conditions were used for this analysis and results reported in multiple TRs, from 3^rd^ TR (around 1.96 sec) to 9^th^ TR (around 5.89 sec). Older adults recruited these areas more than their younger counterparts for all conditions. For visualisation purposes, bootstrap ratio threshold is set at 2 (*p* < 0.05); however, reported whole-brain activity is set at BSR 3 (*p* < 0.001). Abbreviations: L = left hemisphere, R = right hemisphere.

#### Brain-behavior results

##### ACC and rejection rate

Our brain-behavior analysis on ACC and rejection rate revealed one significant latent variable accounted for 15% of covariance of the data (*p* = 0.001). This network comprised of regions such as right medial cingulate cortex, left paracentral lobule, right inferior frontal gyrus, bilateral superior temporal gyrus, left middle temporal gyrus, bilateral middle frontal gyrus, right lingual gyrus, bilateral thalamus, and right hippocampus (Table 4). This network was engaged for believable inferences (believable premise/believable conclusion (BB)) and contributed to more rejection rate in this condition. In other words, individuals who recruited this network more, rejected the believable syllogisms to a greater extent (Figure 3). However, during unbelievable inferences, this network was not engaged among older adults and was not reliably activated for any of conditions among younger adults (CIs crossing zero).

**Table 4.** Functional brain networks of IFG and ACC for all conditions during the decision-making stage.

| Regions | Hem | MNI coordinates | BSR |
| --- | --- | --- | --- |
|  |  | XYZ |  |
| *Anterior cingulate cortex network* |  |  |  |
| Anterior cingulate cortex | R | -10 36 18 | -4.61 |
|  | L | 8 40 4 | -4.84 |
| Medial cingulate cortex | R | 4 -16 38 | -5.83 |
| Paracentral lobule | L | -6 -36 60 | -6.91 |
| Inferior frontal gyrus (p. Orbitalis) | R | 46 18 -12 | -5.90 |
| Inferior frontal gyrus (p. Triangularis) | R | 56 16 22 | -4.68 |
| Middle frontal gyrus | R | 42 34 18 | -5.10 |
|  | L | 40 18 56 | -4.53 |
| Superior frontal gyrus | R | 20 16 60 | -4.95 |
| Superior medial gyrus | L | -8 48 24 | -4.82 |
| Superior temporal gyrus | R | 54 -24 4 | -5.54 |
|  | L | -56 -8 0 | -5.15 |
| Middle temporal gyrus | L | -44 -74 20 | -4.98 |
| Lingual gyrus | R | 22 -44 -2 | -4.83 |
| Cerebellum | R | 8 -74 -14 | -3.97 |
| Cuneus | L | -12 -58 24 | -3.41 |
| Thalamus | R | 26 -20 -2 | -6.18 |
|  | L | -10 -32 6 | -3.54 |
| Hippocampus | R | 36 -18 -16 | -3.84 |
| *Inferior frontal gyrus network* |  |  |  |
| Inferior frontal gyrus (p. Triangularis) | R | 46 2 8 | -16.14 |
|  | L | -50 20 16 | -14.26 |
| Anterior cingulate cortex | R | 18 42 4 | -4.71 |
| Middle temporal gyrus | L | -56 6 -22 | -5.95 |
| Temporal pole | L | -46 8 -22 | -6.80 |
| Inferior parietal lobule | R | -50 -50 38 | -5.54 |
| Middle occipital gyrus | R | 38 -84 0 | -5.48 |
| Cerebellum | L | -20 -50 -16 | -5.32 |
| Precuneus | R | 2 -62 62 | -5.15 |
| Hippocampus | R | 38 -24 -14 | -5.04 |
|  | L | -31 -8 -17 | 4.76 |

Hem = hemisphere; BSR = bootstrap ratio; MNI = Montreal Neurological Institute. Anterior cingulate and inferior frontal gyrus networks were examined within both age groups for all experimental conditions, except neutral condition, including the behavioral performance. Onsets of syllogisms during the decision making, conclusion stage, were included in the analyses.

**Figure 3.**
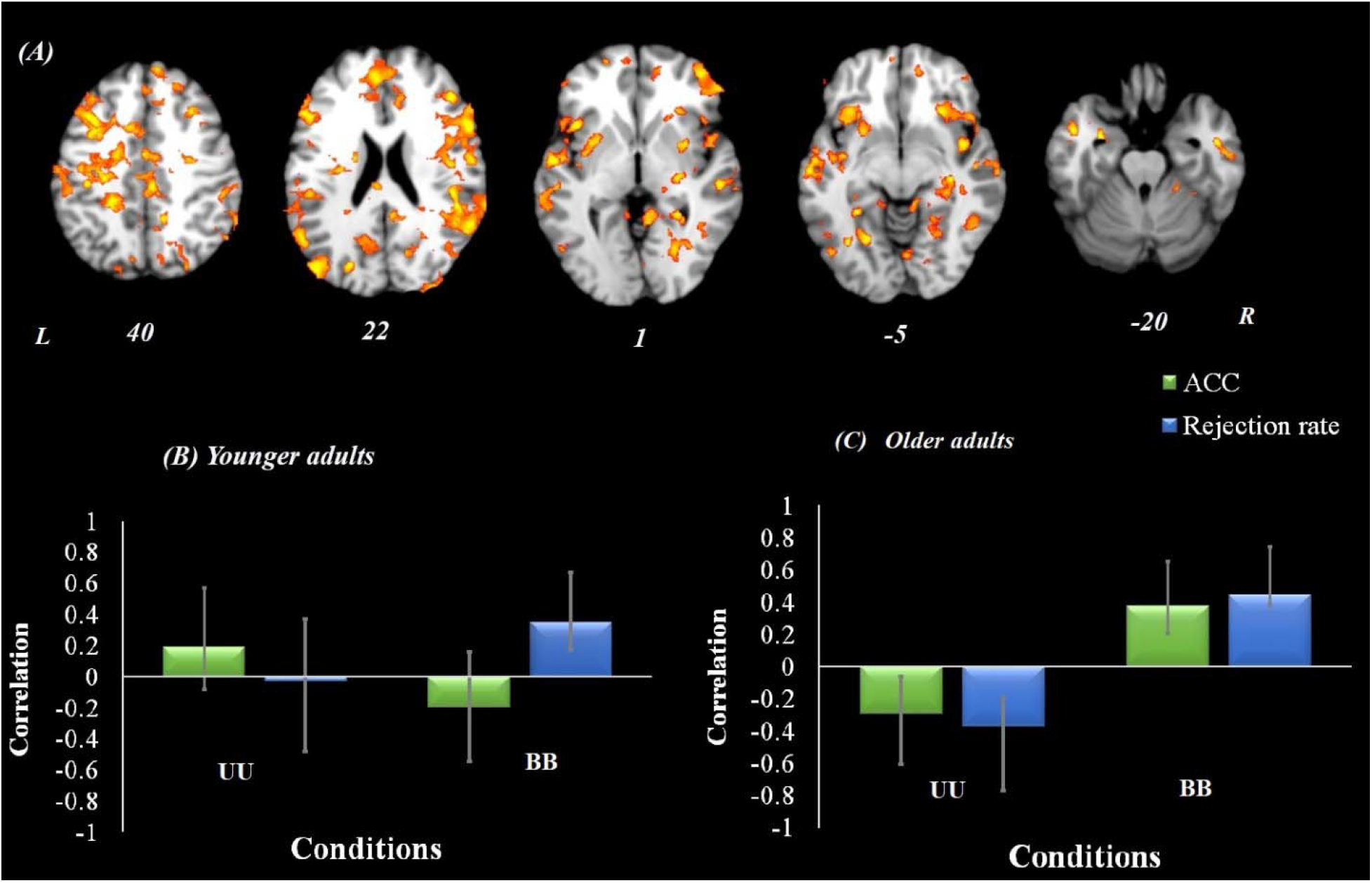
Anterior cingulate cortex functional connectivity results among both age groups. Onsets of conclusion stage were used for this analysis. ***Panel A***, Brain regions shown in yellow were positively correlated with the anterior cingulate and contributed to more rejection for believable inferences (believable premise/believable conclusion; BB) among older adults (Panel C), but none of the effect is significant among younger adults (Panel B). ***Panel B***, A correlation between the brain activity pattern in panel A and behavioral performance among younger adults. ***Panel C***, A correlation between brain activity pattern in panel A and behavioral performance among older adults. All experimental conditions were included in this analysis but for the simplicity of representation, only significant conditions are shown. Peak coordinate used for ACC = [0 32 22]. For visualization purposes, bootstrap ratio threshold is set at 2 (*p* < 0.05); however, reported whole-brain activity is set at BSR 3 (*p* < 0.001). Error bars represent 95% confidence intervals. Abbreviations: ACC = anterior cingulate cortex, L = left hemisphere, R = right hemisphere.

##### IFG and rejection rate

The brain-behavior analysis with bilateral IFG and rejection rate revealed one significant latent variable which accounted for 20% of the covariance of the data (*p* = 0.001). This network included bilateral IFG, right anterior cingulate cortex, left middle temporal gyrus, left temporal pole, right inferior parietal lobule, middle occipital gyrus, and bilateral hippocampus (Table 4). This network contributed to more rejection rate for believable inferences (believable premise/believable conclusion (BB)) and lower rejection rate for unbelievable inferences (unbelievable premise/unbelievable conclusion (UU); Figure 4). This network was not engaged by younger adults, nor contributed to their performance in any of the condition (CIs crossing zero).

**Figure 4.**
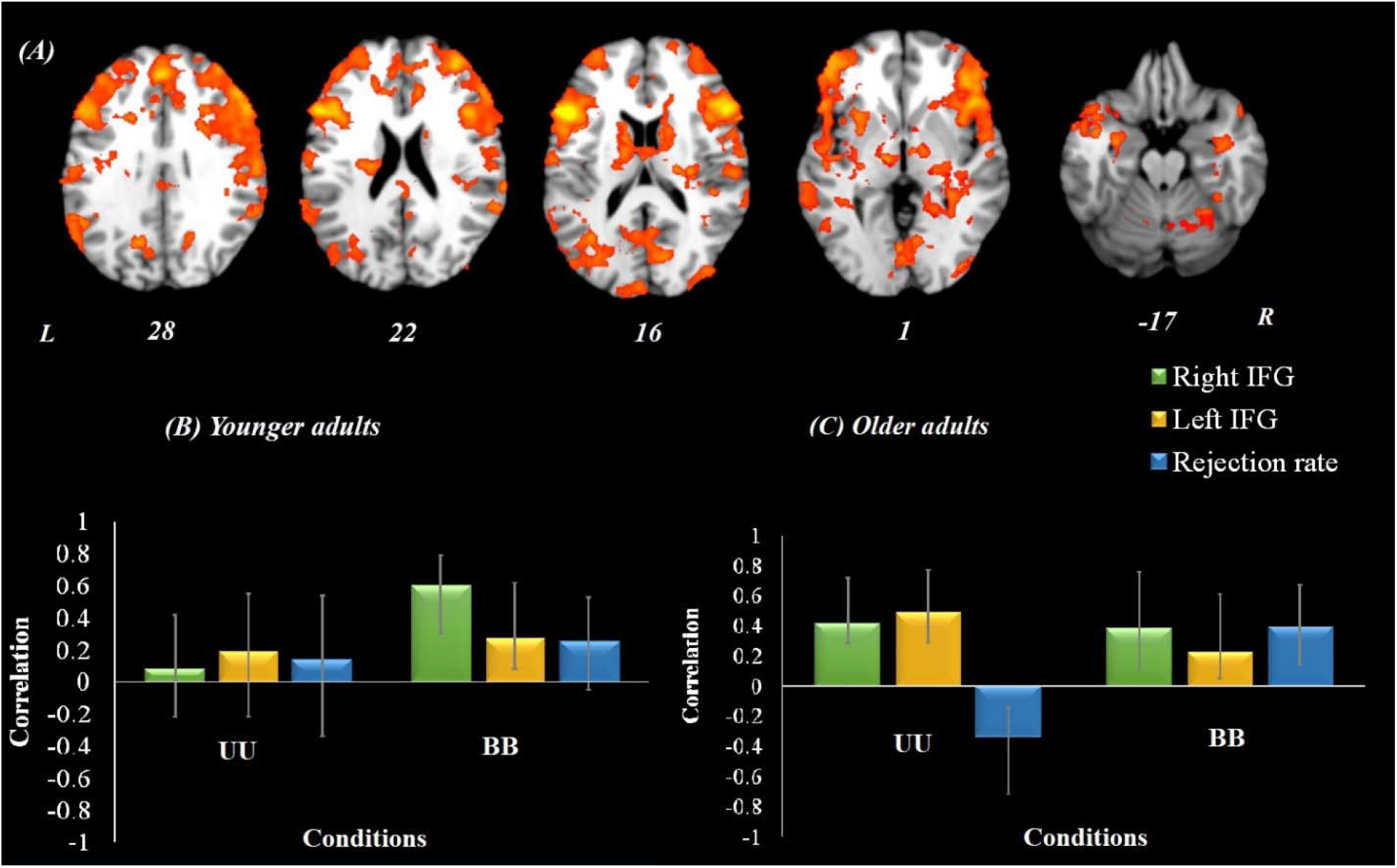
Inferior frontal gyrus functional connectivity results among both age groups. Onsets of the conclusion stage were used for this analysis. ***Panel A***, Brain regions shown in yellow were positively correlated with the inferior frontal gyrus and contributed to more rejection for believable inferences (believable premise/believable conclusion; BB) and less rejection of unbelievable inferences (unbelievalbe premise/unbelievable conclusion; UU) among older adults (Panel C), but none of the effect is significant among younger adults (Panel B). ***Panel B***, A correlation between the brain activity pattern in panel A and behavioral performance among younger adults. ***Panel C***, A correlation between the brain activity pattern in panel A and behavioral performance among older adults. All experimental conditions were included in this analysis but for the simplicity of representation, only significant conditions are shown. Peak coordinate used for IFG = [48 22 8] and [-48 22 16]. For visualization purposes, bootstrap ratio threshold is set at 2 (*p* < 0.05); however, reported whole-brain activity is set at BSR 3 (*p* < 0.001). Error bars represent 95% confidence intervals. Abbreviations: IFG = inferior frontal gyrus, L = left hemisphere, R = right hemisphere, UU = unbelievable premise/unbelievable conclusion, BB = believable premise/believable conclusion.

In sum, the results from the functional connectivity analyses yielded that the IFG network contributed to the rejection rate for both believable and unbelievable inferences, whereas the ACC network was involved in rejection of believable inferences only.

#### Structure-function-behavior results

##### Cingulum bundle and rejection rate

Our structure-function-behavior analysis using the cingulum FA values with rejection rate in both age groups revealed one significant latent variable. This latent variable accounted for 14% covariance of the data (*p* < 0.001) and included regions such as anterior cingulate cortex, bilateral insula, right inferior frontal gyrus, bilateral superior temporal gyrus, bilateral posterior cingulate cortex, bilateral putamen, left hippocampus, and right parahippocampus. This functional network which was correlated with the structural integrity of the cingulum FA values was more engaged during believable inferences (believable premise/believable conclusion (BB)). In other words, individuals with higher structural integrity in the left cingulum tract activated this network more during believable statements. Further analyses using uncinate fasciculus and inferior longitudinal fasciculus did not reveal any significant findings with behavior and BOLD signal.

#### Structure-behavior results

##### Uncinate fasciculus and rejection rate

Next, we examined whether the relation between age group and the rejection rate is mediated by the integrity of the fractional anisotropy (FA) values obtained from these tracts. The results indicated that the relationship between age group and rejection rate was mediated by the integrity of the left uncinate fasciculus tract during believable conclusion, but not unbelievable ones. The total effect of age group on rejection rate was significant (total effect = −0.0041, standard error = 0.0012, *p* < 0.001, Figure 5A, pathway c), and the direct effect was also significant (direct effect = −0.0039, standard error = 0.001, *p* < 0.001). The standardized indirect effect using bootstrapped procedure found to be significant (indirect effect lower 95% CI = −0.005, upper 95% CI = - 0.0001, pathway c’). Older adults’ participants showed lower integrity in the left uncinate fasciculus (Figure 5A, pathway a) and that lower integrity of the left uncinate fasciculus predicted less rejecting of the believable conclusions after controlling for age (Figure 5A, pathway b). The mediation analysis with the right uncinate fasciculus was not significant. This result suggests that the relationship between age and logical reasoning performance during believable statement was mediated by the left uncinate fasciculus integrity. None of the other effects for cingulum or inferior longitudinal fasciculus were significant for mediation analyses.

**Figure 5.**
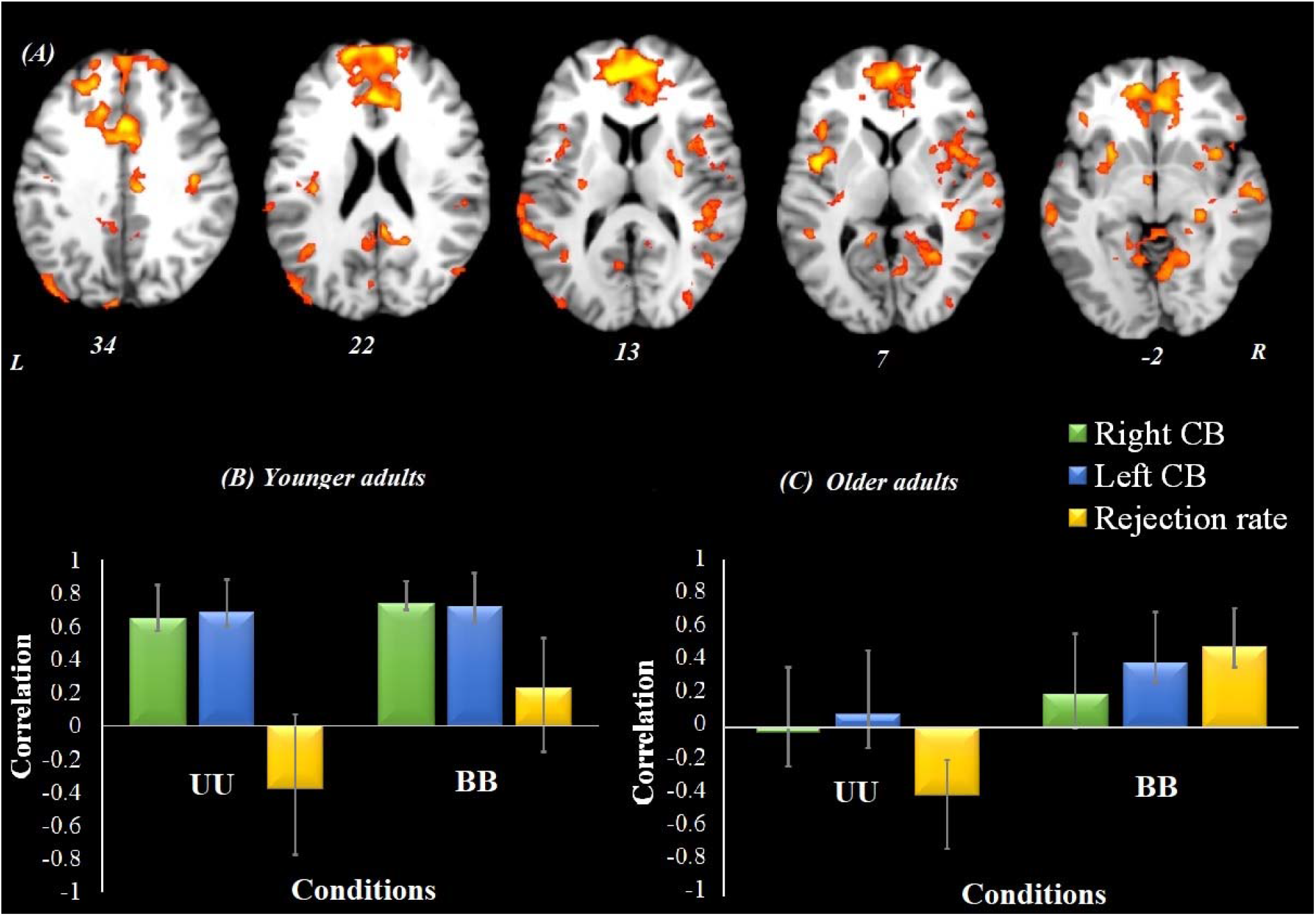
Structure-function-behavior results with cingulum bundle fractional anisotropy and rejection rate. Fractional anisotropy (FA) values of the cingulum bundle (CB) for both left and right sides and rejection rate were included in this analysis. ***Panel A***, Brain regions shown in yellow were positively correlated with the CB FA values and contributed to more rejection for believable inferences (believable premise/believable conclusion; BB) among older adults (Panel C), but not for the unbelievable inferences (unbelievable premise/unbelievable conclusion; UU). This brain network was also correlated with the CB FA values in both conditions among younger adults but did not contribute to behavioral performance, confidence intervals crossing zero. ***Panel B***, A correlation between the brain activity pattern in panel A and CB FA values and behavioral performance among younger adults. ***Panel C***, A correlation between the brain activity pattern in panel A and CB FA values and behavioral performance among older adults. All experimental conditions were included in this analysis but for the simplicity of representation, only significant conditions are shown. For visualization purposes, bootstrap ratio threshold is set at 2 (p < 0.05); however, reported whole-brain activity is set at BSR 3 (p < 0.001). Error bars represent 95% confidence intervals. Abbreviations: CB = cingulum bundle, L = left hemisphere, R = right hemisphere, UU = unbelievable premise/unbelievable conclusion, BB = believable premise/believable conclusion.

**Figure 6.**
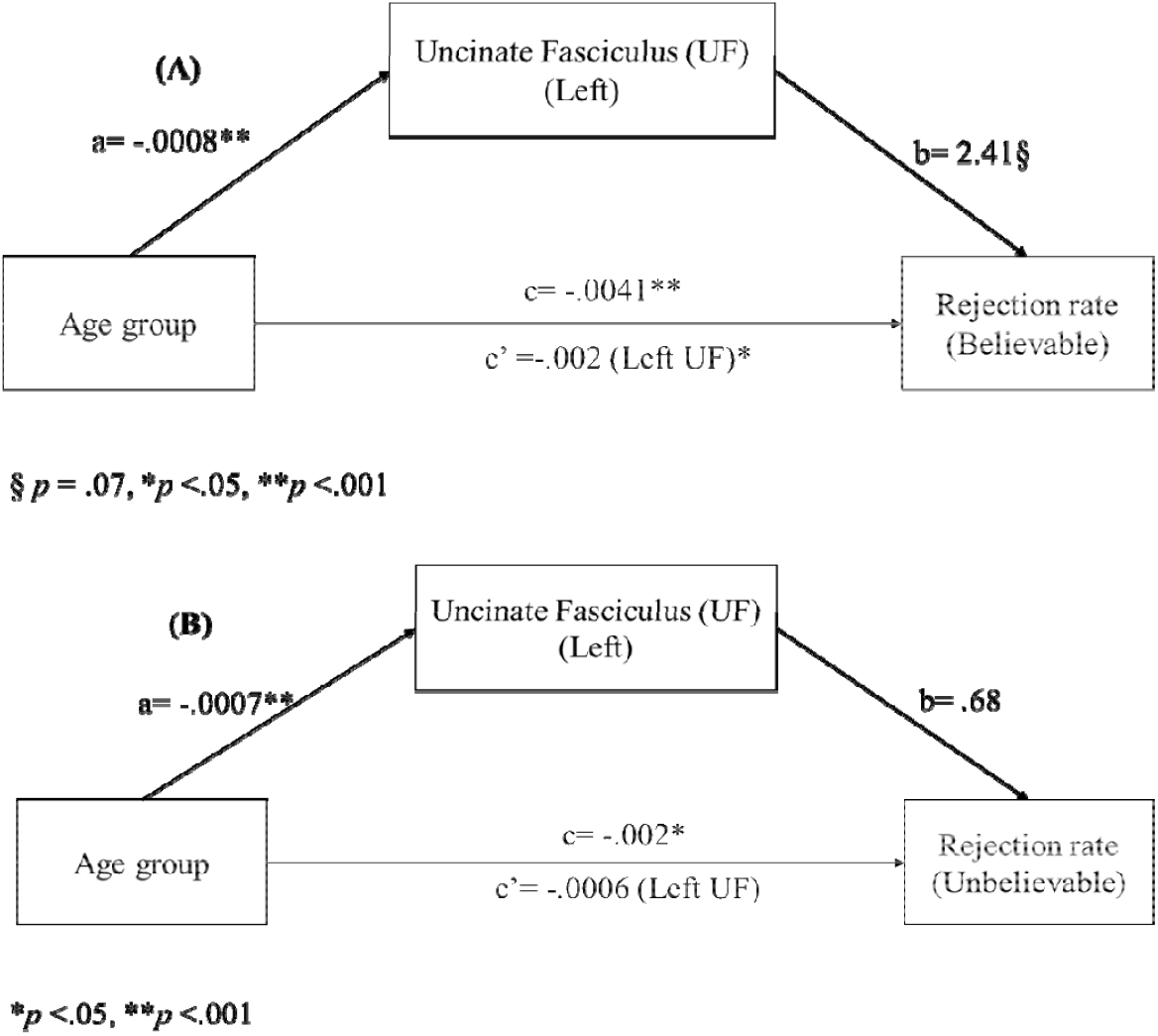
Structure-behavior mediation results with uncinate fasciculus fractional anisotropy (FA) and rejection rate. ***Panel A***, mediation analysis on rejection rate of believable inferences. The numbers along the paths represent standardized regression coefficients. The coefficient below the path from age group to believable inference represents the indirect effect (c’) when left Uncinate Fasciculus FA value was included as a mediator and coefficient above the path represents the direct effect with no mediator in the model. ***Panel B***, mediation analysis on rejection rate of unbelievable inferences. The numbers along the paths represent standardized regression coefficients. The coefficient below the path from age group to believable inference represents the indirect effect (c’) when left Uncinate Fasciculus FA value was included as a mediator and coefficient above the path represents the direct effect with no mediator in the model. Results of mediation analysis with the right uncinate fasciculus was not significant.

## Discussion

The present study provides evidence to explain the age-related differences in logical reasoning and the impact of currently-held beliefs using a syllogism task, supporting inhibitory deficit hypothesis (Hasher & Zacks, 1988). First, the whole-brain task-related functional results showed that older adults engaged a large-scale brain pattern, including anterior cingulate cortex (ACC) and inferior frontal gyrus (IFG), for all conditions more than their younger counterparts during the decision-making stage (conclusion statement). Second, our functional connectivity results suggest that older adults who engaged the ACC network to a larger extent showed better control over their intuitive responses to reject the believable statements (higher rejection rate for believable inferences). Among older adults, the IFG network was also comparably engaged across all conditions during decision-making stage and contributed to better control over responses, leading to higher rejection of believable and lower rejection of unbelievable inferences. Finally, structure-function-behavior analyses showed that the integrity of left cingulum bundle tract and uncinate fasciculus contributed to rejection rate for believable inferences, highlighting the importance of inhibitory control over presumably emotional responses required during logical reasoning task.

### The role of ACC

During the decision-making stage, older adults engaged a large-scale brain pattern including bilateral IFG and ACC for all conditions. We observed sustained engagement of the ACC during the conclusion stage, from earlier stage until the end of the decision-making stage (Figure 4). The ACC – a central region of the executive attention network – has been linked to a variety of self-regulation functions such as monitoring of conflict (Botvinick et al., 2001; Botvinick et al., 2004), regulating emotions (Buhle et al., 2014; Bush et al., 2000; Ziaei et al., 2017), responding to errors (Holroyd & Coles, 2002), and responses-outcome association (Alexander & Brown, 2010), to name a few. In aging, studies have reported a link between ACC activity and performance in a number of tasks such as Stroop (Milham et al., 2002), memory encoding of negative stimuli (Ziaei et al., 2017), and theory of mind (Ziaei et al., 2016). In the context of our findings, engagement of the ACC from the beginning of the conclusion stage emphasizes the importance of engagement of inhibitory control and response monitoring during logical reasoning in earlier stage of decision making. Our functional connectivity results further support this claim and highlight the role of fronto-parietal network functionally connected to the ACC during the task. Critically, the fronto-parietal network contributes to rejecting believable inferences. That is, the more the fronto-parietal network is engaged, older adults are more able to correctly reject believable statements. It has to be noted that rejecting a believable inference that is in line with currently-held beliefs is challenging and requires more cognitive effort. The engagement of fronto-parietal network in deductive reasoning is in line with previous findings (Monti et al., 2009; Reverberi et al., 2007). Our results take these findings further and suggest that the logical reasoning task was cognitively demanding for older individuals, evidenced by behavioral and neuroimaging findings, which resulted in engagement of the fronto-parietal network relative to their younger counterparts.

### The role of IFG

The importance of the IFG has been highlighted for reasoning in previous studies using functional MRI (Prado et al., 2011; Reverberi et al., 2012; Reverberi et al., 2010) as well as repetitive transcranial magnetic stimulation (Tsujii et al., 2011). For instance, Tsujii and colleagues (2011) examined the role of the superior parietal lobe and the IFG during a logical reasoning task in which the believability load of inferences was manipulated. They found that stimulation of bilateral parietal lobe disrupted performance on incongruent – unbelievable – inferences whereas stimulation of the left IFG impaired performance during congruent – believable – inferences and facilitated performance during incongruent – unbelievable – condition. Findings from our functional connectivity results yield a link between the bilateral IFG network, higher rejection of believable, and lower rejection of unbelievable inferences. The increased activity in the network connected to the IFG among older adults might also suggest the cognitive effort exerted during the task. This speculation is supported by the coupling between ACC, parietal lobe, and the IFG, suggesting the cognitive effort needed to perform the task (Berry et al., 2017). While IFG is critical in inhibitory control (Aron et al., 2014), previous studies have also shown that the IFG might be involved in detecting cues regardless of whether the inhibitory control is required (Hampshire et al., 2010). In the context of our task, it is possible that during a logical reasoning task, believability load of the statements might have triggered orientation of attention toward cues that are necessary to initiate inhibitory control over currently-held beliefs.

While most studies agree on the role of IFG in reasoning, nonetheless there are still inconsistencies between findings about the importance of laterality of IFG in various logical reasoning tasks. It seems that a belief-based heuristic system activates the left IFG (Ragni et al., 2016) and inhibition of the default heuristic system (or enabling the logical system) engages the right IFG (Reverberi et al., 2010; Tsujii et al., 2010; Tsujii et al., 2011). The bilateral engagement of the IFG in this study, however, can be due to the use of multivariate method in this study, the PLS. As previously mentioned, the PLS is more sensitive than traditional GLM to detect patterns of activities associated with multiple experimental conditions simultaneously. Given that most previous studies have used univariate method to examine neural correlates of logical reasoning, the bilateral engagement of the IFG in this study demonstrates a methodological advantage of multivariate analysis for examining higher-order cognitive functions such as logical reasoning.

Altogether, our functional connectivity results revealed that sub-regions of the prefrontal cortex, the ACC and IFG, might have slightly different roles for the inhibition of currently-held beliefs. The ACC and its network contributed to currently rejecting believable statements more than other conditions. On the other hand, the IFG and its network contributed to the higher rejection of believable and lower rejection of unbelievable statements. More work is needed to explore the role of other frontal and subcortical brain areas, such as the hippocampus (see Ziaei et al., submitted) during a reasoning task among both age groups and the inhibitory control that is required to outperform such task.

### Structural integrity of cingulum and uncinate fasciculus

Another important finding from the current study was the relationship between cingulum bundle and the rejection rate of believable inferences. Cingulum bundle has been shown to be involved in higher-order cognitive tasks such as attention, inhibitory control (Li et al., 2018; Metzler-Baddeley et al., 2012), executive functioning (Grieve et al., 2007), emotional processing, as well as belief updating (Moutsiana et al., 2015). In the context of aging, it has been shown that that older adults with more integrity in the cingulum tract, would perform differently in the cognitive control tasks than younger adults (Metzler-Baddeley et al., 2012), highlighting the importance of cingulum integrity in inhibitory control. Because the cingulum relates to medial and dorsal prefrontal cortex, its role in inhibitory control has been repeatedly shown in the literature. Therefore, given the importance of inhibitory control in logical reasoning, it is not surprising that individuals with more integrity in this tract would be able to inhibit their currently-held beliefs and thus, perform better during believable conditions. As expected, rejecting believable statements require higher inhibitory control and thus our structure-function-behavior results are in line with our prediction that individuals with more integrity in the cingulum tract can inhibit their intuitive/heuristic tendency and reject believable statement more. Additionally, our results are in line with the functional connectivity results once again supporting the anterior cingulate’s role in inhibition of response during believable conditions.

We also found a relationship between uncinate fasciculus and the rejection rate for believable inferences. The uncinate fasciculus has been implicated in a number of functions, such as language, memory, and social-emotional processing (Von Der Heide et al., 2013). This tract is usually associated with the limbic system and connects the prefrontal cortex, including Broadman area 10, 9, to the temporal lobes through bidirectional pathways (Hasan et al., 2009). This tract is often associated with the limbic structure and is involved in emotion and memory (Olson et al., 2015; Von Der Heide et al., 2013), emotional empathy (Oishi et al., 2015), and in diseases with the hallmark of temporal lobe functional deficiency such as depression (Zhang et al., 2012), epilepsy (Von Der Heide et al., 2013), and mild cognitive impairement and Alzheimer’s disease (Fouquet et al., 2009; Rémy et al., 2015; Villain et al., 2010). The lower integrity of the left uncinate fasciculus contributed to the lower rejection of believable statements. This finding is in line with previous results on updating beliefs that have reported activation of frontal lobes and subcortical areas such as hippocampus and amygdala (Li et al.; Moutsiana et al., 2015). It is not surprising that the limbic area would be engaged during a logical reasoning performance. Specifically, the hippocampus is expected to be involved in semantic retrieval of currently-held assumptions, and the amygdala’s activity is expected to be elicited by the emotional response depending on the belief load. Thus, the better integrity of the uncinate fasciculus tract indicates better ability to retrieve previously stored information as well as better control over emotional responses, which could result in inhibition of beliefs and better logical reasoning performance overall. Our results indicate that older adults’ deficit in cognitive control ability during the logical reasoning performance might stem from a lower integrity in white matter structures, such as cingulum bundle and uncinate fasciculus. So that older adults with preserved integrity in these tracts would be able to control their intuitive responses better and to reject the believable statements when necessary.

### Limitations

One of the limitation of this study is the lack of comparison between various logical reasoning tasks such as inductive or probabilistic reasoning tasks. The use of syllogistic reasoning allowed manipulation of belief loads and comparison between stages of reasoning. However, future studies are needed to investigate age-related differences in functional and underlying structural brain networks in relation to behavioral performance during different types of logical reasoning tasks. Additionally, while both age groups performed highly in the task, older participants, particularly, were high performers. Although the performance difference between the two age groups was statistically significant, with older adults’ performance was lower than younger adults specifically for the unbelievable statements, the average performance difference between the two groups was not substantial (the standard errors overlapping). Therefore, the current behavioral results need to be considered cautiously and future work is needed to replicate these findings among older adults with different levels of cognitive capacities and with syllogisms in which their difficulty levels are manipulated to elicit belief bias (Brisson et al., 2014).

### Conclusion

The primary aim of this study was to examine age-related differences in functional networks of ACC and IFG during a logical reasoning task and to investigate if these networks are modulated by the believability load of the inferences among two age groups. Our whole-brain task-related and functional connectivity results support the critical role of inhibitory control and response inhibition – reflected in the functional networks from ACC and IFG – during logical reasoning among older adults. Interestingly, through analysis of the relationship between structural integrity of white matter tracts and behavioral performance, our results further support that the structural integrity of these tracts and their connections are underlying the logical reasoning performance. In summary, our behavioral, functional, and structural data converging that older adults have more difficulties in inhibition of their current beliefs reflected in the engagement of wide-spread fronto-parietal networks during the syllogistic reasoning task. Furthermore, our results provide a possible explanation suggesting that the white matter integrity of specific tracts might be an underlying factor contributing to the age-related differences in inhibitory control over currently-held beliefs.

Difficulties in reasoning and drawing conclusions from assumptions might have serious consequences in many instances, ranging from simple and personal instances to complex and political ones. Incongruency of given assumptions with currently-held personal beliefs in different circumstances adds to the complexity and the importance of this topic. It seems that our ability to inhibit currently-held beliefs, required for a well-construed conclusion, changes with age. These findings are well aligned with previous studies suggesting that prior knowledge disrupts learning and subsequent performance among older adults (for a review see Spreng & Turner, 2019). These findings have important and broader implications for situations where inhibitory control is compromised, due to fatigue or drug use, and as a result decision making would be heavily affected, especially if individuals are emotionally involved.

## Acknowledgment

The authors would like to thank our participants for their time and acknowledge the practical support provided by the imaging staff at the Centre for Advanced Imaging. We would also like to thank Mr. Graham Phegan for his time and help in editing the manuscript and Ms. Ashley York for her help with the data collection.

## Compliance with ethical standards

### Funding

This work was supported by funding from the Australian Research Council Science of Learning Special Research Initiative (SR120300015) to DCR.

### Ethical approval

All procedures performed in studies involving human participants were in accordance with the ethical standards of the national research committee and with the 1964 Helsinki declaration and its later amendments or comparable ethical standards.

### Informed consent

Informed consent was obtained from all individual participants included in this study.

